# Genome-resolved metagenomics and detailed geochemical speciation analyses yield new insights into microbial mercury cycling in geothermal springs

**DOI:** 10.1101/2020.02.03.933291

**Authors:** Caitlin M. Gionfriddo, Matthew B. Stott, Jean F. Power, Jacob M. Ogorek, David P. Krabbenhoft, Ryan Wick, Kathryn Holt, Lin-Xing Chen, Brian C. Thomas, Jillian F. Banfield, John W. Moreau

## Abstract

Geothermal systems emit substantial amounts of aqueous, gaseous and methylated mercury, but little is known about microbial influences on mercury speciation. Here we report results from genome-resolved metagenomics and mercury speciation analysis of acid warm springs in the Ngawha Geothermal Field (<55 °C, pH < 4.5), Northland Region, Aotearoa (New Zealand). Our aim was to identify the microorganisms genetically equipped for mercury methylation, demethylation, or Hg(II) reduction to volatile Hg(0) in these springs. Dissolved total and methylated mercury concentrations in two adjacent springs with different mercury speciation ranked among the highest reported from natural sources (250–16000 ng L^−1^ and 0.5–13.9 ng L^−1^, respectively). Total solid mercury concentrations in spring sediments ranged from 1273 to 7000 µg g^−1^. In the context of such ultra-high mercury levels, the geothermal microbiome was unexpectedly diverse, and dominated by acidophilic and mesophilic sulfur- and iron-cycling bacteria, mercury- and arsenic-resistant bacteria, and thermophilic and acidophilic archaea. Integrating microbiome structure and metagenomic potential with geochemical constraints, we constructed a conceptual model for biogeochemical mercury cycling in geothermal springs. The model includes abiotic and biotic controls on mercury speciation, and illustrates how geothermal mercury cycling may couple to microbial community dynamics and sulfur and iron biogeochemistry.

**IMPORTANCE:** Little is currently known about biogeochemical mercury cycling in geothermal systems. This manuscript presents an important new conceptual model, supported by genome-resolved metagenomic analysis and detailed geochemical measurements. This work provides a framework for studying natural geothermal mercury emissions globally. Specifically, our findings have implications for mercury speciation in wastewaters from geothermal power plants and the potential environmental impacts of microbially and abiotically formed mercury species, particularly where mobilized in spring waters that mix with surface- or ground-waters. Furthermore, in the context of thermophilic origins for microbial mercury volatilisation, this report yields new insights into how such processes may have evolved alongside microbial mercury methylation/demethylation, and the environmental constraints imposed by the geochemistry and mineralogy of geothermal systems.

## Introduction

Geothermal springs and fumaroles emit substantial amounts of aqueous and gaseous mercury (Hg) (1). Aqueous Hg concentrations in these systems often exceed 100 ng L^−1^ and total Hg levels can approach 25 µg L^−1^ (2–4). Despite these ultra-high mercury levels, few studies have examined biotic and abiotic mechanisms for Hg transformations or Hg speciation in geothermal springs. Specifically, the potential for native thermophiles to mediate mercury transformations, i.e., reduction of Hg(II) to Hg(0) or methylation/demethylation of mercury (to CH_3_Hg^+^ or “MeHg”; or to Hg[II], respectively; (5–7) remains poorly understood.

Hg species have a high binding affinity to thiols and lipids, causing damage to proteins, enzymes, and nucleic acids. They can inhibit microorganisms at sub-micromolar concentrations (8). Microorganisms living in environments with elevated Hg (>100 ng L^−1^) commonly possess genes encoding for reduction of Hg(II) to volatile Hg(0), as a means of Hg detoxification (9, 10). The *mer* operon is used by many bacteria and archaea to detoxify Hg(II), by converting it to volatile Hg(0) (7, 10, 11). Interestingly, *mer* has a phylogenetic origin in the thermophiles (12, 13). Additionally, microbes equipped with the organomercurial lyase, encoding *merB* gene as part of the *mer*-operon, are able to detoxify organic Hg compounds, in tandem with mercuric reductase (MerA), to produce methane (CH_4_) and Hg(0) (9, 14). Finally, some anaerobic bacteria are suspected to be able to oxidatively demethylate Hg species independent of the *mer* pathway, producing carbon dioxide (CO_2_) and Hg(II) (15), although a biochemical pathway has not been identified for *mer*-independent demethylation.

The efficiency of microbial methylation of Hg(II) to MeHg is still largely unknown in geothermal spring ecosystems, particularly under acidic conditions (<55 °C, pH < 4.5; (4, 5, 16)). However, the *hgcAB* genes required by microorganisms to methylate Hg (17) have been reported from many environments, including wetland sediments, rice paddy soils, thawing permafrost, hypersaline and hypersulfidic waters, soda lakes, and geothermal systems (16, 18–20). These environments typically host abundant sulfate- and iron-reducing bacteria (Deltaproteobacteria), as well as methanogenic and acetogenic Methanomicrobia (Euryarchaeota), Chloroflexi, and Firmicutes, all of which contain species capable of Hg methylation (18, 21–23). Metagenomic analyses have also identified *hgcAB* genes in Chrysiogenetes, Atribacteria (candidate phylum OP9), and candidate phylum ACD79 (16). Although Hg methylation is most often associated with sulfate reduction, the existence of environmental triggers or controls on Hg methylation remains poorly understood, as does the evolution and phylogenetic distribution of *hgcAB* genes.

By comparison, in non-geothermal environments, elevated Hg concentrations can inhibit microbial Hg methylation (24), and lead to conditions favoring MeHg demethylation (25) or methylation-demethylation cycles (26–28). However, in-depth understanding of Hg methylation in geothermal springs, similarly to acid mine drainage (AMD) (29), has not been established. Conversely, acidophiles in AMD systems have been well studied with respect to potential for metal and sulfur cycling (30), but with less focus on Hg speciation and methylation. Here, we combined metagenomic and geochemical speciation analyses to understand Hg transformations in the context of biogeochemical cycling (e.g., S, Fe) in an acidic warm spring microbiome. In order to understand physicochemical constraints on microbial Hg transformations, we studied Hg speciation across a gradient of environmental factors that can influence microbiome composition and activity. We compare our findings to those of previous geothermal spring studies, to refine the conceptual model for geothermal Hg cycling.

## Materials and Methods

### Site description

The Ngawha Geothermal Field (NGF) constituted the field site for investigating Hg cycling in a low pH (<4.5), elevated Hg (>100 ng L^−1^), and sulfide-rich (>0.1 mg L^−1^) environment (Figure S1). Mercury ore deposits in the NGF occur as cinnabar, metacinnabar and native Hg(0) in association with active hot springs, fumaroles, and mud pools (31). Elemental Hg (mainly gaseous Hg(0)), travels from deep geological sources to the surface, either in hydrothermal fluids or geothermal gases, where it reacts with oxygen in presence of chloride to form Hg(II); Hg(II) in turn reacts with dissolved sulfide (biogenic) to precipitate cinnabar (32). Roughly 33,000 kg of cinnabar ore was mined from the Tiger Springs area at Ngawha during the first half of the 20th century (31). Tiger Springs and other areas within the NGF still host an active geothermal system that releases approximately 530 kg of Hg_T_ annually, ~44% of which is thought to be emitted to the atmosphere (32). The remaining Hg resides in the local surficial waters and sediments (32). Cinnabar precipitation was confirmed by X-ray diffraction analysis of Tiger Springs sediments and nearby topsoil (Fig. S2).

### Sampling techniques

Several springs from three areas, Tiger Springs (TS), Ginn Ngawha Spa (GN), and Ngawha Springs Baths (NS), were sampled in April and October of 2011. Many of the springs were edged with boards for use as soaking baths. Water samples for Hg analyses were filtered through 0.45 μm membrane syringe filters, preserved with 1% v/v reagent grade HCl, and stored in acid-washed HDPE bottles in the dark. Filtered water samples (0.45 μm) for anion analysis were stored in sterile 50 mL plastic Falcon tubes. All filtered samples were stored at 4°C. Redox potential (mV) and pH measurements were taken at each sampling site using an Orion Model 250A portable meter with a glass pH electrode; spring water temperature measurements were also taken at the time of sample collection. Sediment samples were collected from the floor or wall of each spring, as well as from bulk water samples, into sterile 50 mL Falcon tubes. Using sediments collected on the bottom of the boarded spring bath and suspended in water allowed us to capture a mix of aerobic and anaerobic members of the microbial community, as well as those that live across a range of temperature gradient from the cooler waters (<45 °C) to the hotter sediments (>55 °C). Samples were stored on ice for ~4 days until they could be transferred to laboratory storage at −80 °C (i.e. transported from NZ to Melbourne, Australia). Sediment samples were used for Hg_T_ analyses and for whole community DNA extractions.

### Hg and MeHg analyses

Total Hg and MeHg concentrations of filtered waters and freeze-dried sediments collected in April and October 2011 were measured at the Wisconsin Water Science Center, (WWSC, U.S. Geological Survey, Middleton, Wisconsin) on a Perkin-Elmer Elan 9000 quadrupole ICP-MS and a Brooks Rand Atomic Fluorescence Spectrophotometer Model III, respectively. Filtered water samples were analyzed within six months of sampling at the Wisconsin Mercury Research Lab (WMRL) of the Wisconsin Water Science Center (USGS; Middleton, Wisconsin). Filtered water samples analyzed for Hg_T_ species were treated with BrCl solution to ensure all Hg species in the sample were oxidized to Hg(II). Prior to analysis, SnCl_2_ solution was added to the vials which reduced the Hg(II) species to volatile Hg(0). The samples were then ethylated, purged with argon gas, and analyzed by GC (using Brooks Rand Autosampler and Total-Hg Purge and Trap system) in tandem with atomic fluorescence spectrometry. Methylmercury analysis of filtered waters was determined by distillation, gas chromatography separation, and speciated isotope dilution mass spectrometry using ICP-MS, following USGS Method 01-445 and WMRL standard protocols (33). Four blanks and two duplicate spikes were included in each run for quality assurance. Method detection limits for total and methylated mercury were 0.007 ng, and 0.03–1.2 ng L^−1^, respectively (depending on the dilution factor required for each sample).

Sediment samples (~5 g wet weight) were freeze-dried overnight on a Heto-Drywinner vacuum system before shipment in sterile glass vials to the WMRL. For solid Hg_T_ analysis, freeze-dried samples were digested with 3:1 HCl:HNO_3_ overnight in a Teflon vessel. The digested sample was then oxidized with BrCl solution. Total Hg analysis was then performed by the same procedure as for filtered water. The detection limit for the solid Hg_T_ analysis was 0.2 ng. Field blanks for Hg analyses were prepared using ultrapure reaction grade water spiked with 1% v/v ultrapure HCl stored in each type of sampling material until analysis. The Hg_T_ and MeHg_T_ values for field blanks were 0.32 ng L^−1^ and 0.57 ng L^−1^, respectively.

### Hg(0) analysis

Vapor samples were collected in October 2011 at a height of 5-10 cm over Tiger and Cub Baths (Tiger Springs area) on SKC Anasorb Sorbent tubes (Model C300). These sorbent tubes are typically used to measure passive exposure to mercury in industrial settings (34, 35). Flow rate (2 L min^−1^). The volume of air sampled was regulated using an SKC Sample Air Pump (PCXR4). Gas samples were passed through a soda lime trap before collection on the hopcalite sorbent to trap excess condensation and neutralize acid. Once used for sampling, sorbent tubes were sealed with Teflon tape and sent to ChemCentre (WA, Australia) to be analyzed per NIOSH Method 6009. The sorbent material was dissolved and oxidized in 1:1 HNO_3_:HCl. This solution was then diluted with DI water, and immediately before analysis on a cold vapor atomic absorption spectrometer (CVAAS), 10% SnCl_2_ was added to reduce all Hg(II) to Hg(0). This Hg(0) was then purged into the CVAAS analyzer (detection limit = 0.01 µg) in an argon gas stream. Sampling of Tiger and Cub Baths was performed under similar conditions. First, a blank Anasorb tube was exposed to ambient levels of Hg (unsealed, with no air pumped through). An additional blank, that remained sealed, was also included in analyses. Blank samples were below the method detection limit (<0.01 µg). Samples taken from the same site, with varying pump times (20-30 minutes) did not yield similar adsorbed Hg concentrations. A positive correlation between total Hg adsorbed versus volume of air pumped through the adsorbent trap (r^2^=0.976, n =4), indicated that saturation of the adsorbent was not reached.

### Common ion analysis

Common anion concentrations (F^−^, Cl^−^, Br^−^, NO_3_^−^, SO_4_^2−^) were measured in unacidified, filtered water samples collected in April 2011, using a Dionex DX-120 ion chromatograph with an IonPAC As14 column (4 × 250 mm) in the Department of Chemistry at The University of Melbourne. Samples were stored in the dark at 4 °C for 4 months between sample collection and analysis. The instrument was set to the following conditions: 1.4 mL min^−1^ flow rate, the eluent was 4.8 mM sodium carbonate and 0.6 mM sodium bicarbonate, with an injection volume of 25 µL. Chromatographs were viewed on PeakNet Run System I software. Method detection limit for IC analyses was 0.0005 mg L^−1^. Determination of sulfide in water samples was performed on filtered water samples prepared in the field for analysis in October 2011 using the Methylene Blue Method (36) and the Hach DR 2800 spectrophotometer (Method 8131). Total Fe was measured in filtered, acidified water samples collected in October 2011 using the Ferrozine method (37), with prepared reagents coordinating to Hach Method 8147. Method detection limit for sulfide measurements was 0.02 mg L^−1^and for Fe measurements was 0.5 mg L^−1^. The samples had been stored in the dark at 4°C prior to analysis. Sample blanks were prepared at time of sample collection, using MQ water.

### DNA extraction

DNA was extracted from sediments collected from Tiger (TS1) and Cub (TS2) Baths and surrounding sites (TS3–TS5) in October 2011 using the MO-BIO PowerMax DNA Isolation Kit, using an alteration to the manufacturer’s protocol. Approximately 5 g of sediment was used for extraction. Sediments were first treated with 5 mL of 10 mM TrisCl (solution C6), the samples were then centrifuged at 6,000 g for 5 minutes and the supernatant was decanted. Then, bead solution C1 was added, and the alternative lysis method in the protocol was followed, which replaces the 10-minute bead-beating step with incubation in a water bath at 65 °C for 30 minutes, followed by vortexing the sample for 1 minute (38). This alternative lysis method was used to reduce shearing of the DNA. Duplicate DNA extractions were performed to ensure a sufficient mass of DNA was extracted for each sample. Replicate DNA extractions were concentrated onto the same spin filter and cleaned using QIAquick PCR purification kit (Qiagen, California).

### Metagenomic analysis

Approximately 113–216 ng of genomic DNA extracted from samples collected in October 2011 (TS1A and TS2A) was barcoded by sample and prepared for sequencing with the Nextera®XT kit, and ten samples were run on a single lane of an Illumina® HiSeq 2500 with 2 × 100 bp paired-end sequencing (Australian Genome Research Facility, Melbourne, AU). This produced ~9 giga-bases of sequence data for two Ngawha metagenomes, Tiger Bath (NW1) and Cub Bath (NW2), out of 43 giga-bases for all ten samples (Table S1). Sequences were binned by bar code, quality filtered, and trimmed to remove Illumina® adapters using Trimmomatic (39). Metagenomes were examined after assembling short reads into longer contigs with IDBA-UD (40). Assembled contigs were uploaded and binned for analysis using ggKbase, and have been made publicly available (http://ggkbase.berkeley.edu/).

### Detection of Hg cycling genes in metagenomes

The metagenomic read sets were screened directly for sequences sharing homology with *hgcA* and *hgcB*, using a hidden Markov model (HMM) method described previously (41). Metagenomic read sets were also screened directly for sequence homology with *merA*, using an HMM-search built previously (41). Assembled metagenomic sequences were searched for *mer*-operon, *hgcA* and *hgcB* genes using ggKbase. Genes were annotated in ggKbase by BLAST searches against the NCBI database (42). Multiple *merA* sequences were extracted from ggKbase, translated to amino-acids using the bacterial translation table, and aligned using ClustalW (43) in MEGA6 (44) with MerA reference sequences from confirmed *mer* operon-possessing microorganisms (13).

### Genomic binning and phylogenetic analyses

Genomic binning of metagenomic data was performed using ggKbase, based on scaffold coverage, GC content of scaffold sequences, and common taxonomy of the contigs. Bins were further refined with emergent self-organizing mapping (ESOM) (45). Scaffold coverage was calculated using Bowtie2 to map reads to the assembled sequences (46). Genome completeness was estimated from the presence of bacterial single copy genes (51 in total) or archaeal single copy genes (38 in total) (Table S2). Several bins, NW1_Thermoplasmata_unknown_1, NW2_Desulfurella_acetivorans_33_49, and NW2_Rhodospirillales_68_7, are likely multi-genome bins that could not be refined to single genomes from coverage and average GC splits. Information regarding genomic bins for each metagenome, as well as unbinned scaffolds, is provided in Table S2. Taxonomy was assigned to the metagenome-assembled genomes based on consensus classification of contigs. There are 34 ribosomal proteins considered universal among bacteria, archaea, and eukaryotes, and which can be used to infer phylogeny (47). In this study, we used ribosomal protein s3, a single-copy gene present in every genomic bin from this study, to compare phylogenies and coverage across each metagenome-assembled genome. Ribosomal protein sequences were pulled from ggKbase and then aligned to translated ribosomal protein s3 sequences from the NCBI non-redundant protein database.

### Hg cycling genes from publicly available hot spring metagenomes

To compare Hg-cycling genes from NGF to those of other geothermal settings, assembled metagenomes from Yellowstone National Park (YNP) were searched for *hgcA* (Table S3) and *merA* genes (Table S4). Included in the phylogenetic analysis were HgcA protein sequences extracted from YNP metagenomic datasets (https://img.jgi.doe.gov) (Table S3). Sequences were obtained by querying all assembled hot spring metagenomic datasets for sequences sharing homology to carbon monoxide dehydrogenase (pfam03599). HgcA proteins share homology with carbon monoxide dehydrogenases and are often mis-annotated as such in microbial genomes (17). HgcA sequences were differentiated from carbon monoxide dehydrogenases using the HMM model described above, with an inclusion value cut-off of 1e-7, and only sequences that included the conserved cap-helix region of HgcA were kept for analyses. Out of a total of 234 hot spring metagenomes (of which 216 were from YNP springs) searched (as of July 2016) there were 4,520 matches to sequences annotated as carbon monoxide dehydrogenases (pfam03599) (2,148 in YNP metagenomes). Of these sequences, 102 were identified as HgcA sequences from 18 different metagenomes (Table S3). Included in the phylogenetic analyses were 65 HgcA sequences found in 15 YNP metagenomes. The *hgcA* genes were found in YNP metagenomes sampled from four sites: Mushroom Springs (Gp0111644–45, Gp0057794), Octopus Springs (Gp0057360, Gp0057796, Gp00111632, Gp0111634, Gp0111638, Gp0111642, Gp0111646), Obsidian Pool (Gp0056876), and Fairy Spring (Gp0051404). Environmental metadata were only provided for a small number of these metagenomes. Sequences from Fairy Geyser Spring (Gp0051404) are from an alkaline (pH: 9–9.2) and mesophilic (33.3–36 °C) phototroph-dominated mat, while sequences from Mushroom Spring (Gp0111644–45, Gp0057794) were from the undermat layer (~3–5 mm) of an anoxygenic and phototrophic microbial mat in an effluent channel of the alkaline bath. The water above the mat had a recorded temperature of 60 °C (48). A separate metagenomic study has reported temperature and pH measurements for Mushroom Spring (60 °C; pH, 8.2), Obsidian Pool (56 °C; pH, 5.7), and Octopus Spring (80 – 82 °C; pH, 7.9) (49). The additional 37 *hgcA* sequences were from assembled metagenomes sequenced from Dewar Creek Spring (77 °C; pH, 8.0) and Larsen North spring, in British Columbia, Canada (50).

## RESULTS

### Water and sediment chemistry

Chemical and physical data for each spring are shown in Tables 1 and 2. Chloride concentrations exceeded 400 mg L^−1^ in all springs except Cub Bath (TS2) at 29.8 mg L^−1^. Across the NGF, sulfate concentrations varied from 9.3 – 1200 mg L^−1^, with the highest concentrations found in springs of pH < 4 (Table 1). In Tiger Springs, Cub Bath had the lowest sulfate concentration (319 mg L^−1^), while the other TS sites were > 1000 mg L^−1^. Sulfide concentrations were nearly 3.5x higher in Cub Bath (6.45 mg L^−1^) than in Tiger Bath (TS1) (1.82 mg L^−1^). The total iron concentration in Tiger Bath (12.7 mg L^−1^) in October 2011 was twice that of Cub Bath (6.06 mg L^−1^).

**Table 1.** Anion and cation measurements in filtered water from hot springs in the Ngawha geothermal field.

| Sample Site | GPS Coordinates (Lat, Long) | F<sup>-</sup><br>mg L<sup>-1</sup> | Cl<sup>-</sup><br>mg L<sup>-1</sup> | Br<sup>-</sup><br>mg/L<sup>-1</sup> | NO<sub>3</sub><sup>-</sup><br>mg L<sup>-1</sup> | SO<sub>4</sub><sup>2-</sup><br>mg L<sup>-1</sup> | S<sup>2-</sup><br>mg L<sup>-1</sup> | Fe<sub>T</sub><br>mg L<sup>-1</sup> |
| --- | --- | --- | --- | --- | --- | --- | --- | --- |
| Tiger Bath (TS1) | -35.4075155°, 173.8554535° | 2.1 | 470 | 2.79 | b.d.l | 1217.8 | 1.82 ± 0.13 | 12.7 |
| Cub Bath (TS2) | -35.4075150°, 173.8554755° | 1.9 | 30 | 0.91 | 0.74 | 319.0 | 6.45 ± 0.15 | 6.06 |
| Hole to right of Tiger Bath (TS3) | -35.4075160°, 173.8554315° | b.d.l | 395 | b.d.l | b.d.l | 1176.9 | 0.87 ± 0.47 | n.m |
| Hole 3m from Tiger Bath (TS4) | -35.4075073°, 173.8554202° | b.d.l | 591 | b.d.l | b.d.l | 1017.0 | n.m | n.m |
| Cinnabar Bath (GN1) | -35.4049290°, 173.8600079° | b.d.l | 876 | b.d.l | b.d.l | 10.2 | n.m | n.m |
| Jubilee Bath (GN2) | -35.4049108°, 173.8600182° | b.d.l | 1000 | b.d.l | b.d.l | 91.9 | n.m | n.m |
| Twin Bath (GN3) | -35.4049380°, 173.8600082° | 2.6 | 1073 | 5.02 | b.d.l | 119.9 | n.m | n.m |
| Kotanitanga (NS1B) | -35.4056976°, 173.8583837° | b.d.l | 661 | b.d.l | b.d.l | 155.1 | n.m | n.m |
| Adjacent to Favourite Bath (NS3B) | -35.4055000°, 173.8583547° | b.d.l | 639 | b.d.l | b.d.l | 192.5 | n.m | n.m |
| Waikato Bath (NS4B) | -35.4055887°, 173.8584129° | b.d.l | 537 | b.d.l | b.d.l | 54.1 | n.m | n.m |

n.m = not measured, b.d.l = below detection limit (0.0005 mg L<sup>-1</sup>)

Total Hg and MeHg_T_ measurements of filtered waters from NGF hot springs sampled in April 2011 are shown in Table 2. In comparison to background levels for freshwater systems, Hg_T_ was relatively high at ~250 – 16,000 ng L^−1^, compared to 0.4 – 74 ng L^−1^ for non-geothermal lakes, and 1 – 7 ng L^−1^ for rivers and streams (51). Non-thermal waters in the Ngawha region are reported to contain 300 – 500 ng L^−1^ Hg (32). Previously reported values for dissolved Hg in NGF thermal waters range from 1,000 – 350,000 ng L^−1^ (32). In this study, the highest levels of Hg_T_ were found in the Tiger Springs region (all > 1000 ng L^−1^) and from Cinnabar Bath (GN1) in the Ginn Ngawha Spa (1460 ng L^−1^) (Fig. S3). Methylmercury levels ranged from 0.5 – 14.0 ng L^−1^; however, elevated concentrations of MeHg_T_ did not correlate with higher Hg_T_ concentrations. Concentrations of Hg_T_ and MeHg_T_ in filtered water samples from NGF were compared to reference values (Table 2). MeHg levels in filtered water samples from YNP were substantially lower than those observed at NGF, with most YNP springs having concentrations below the detection limit (0.013 ng L^−1^). Those with detectable MeHg were still relatively low at 0.026 – 0.080 ng L^−1^ (5). Only three sites sampled in the NGF exhibited a significant proportion of Hg_T_ as MeHg (~1% v/v): Cub Bath (TS2), Kotanitanga Bath (NS1B), and a drainage pool adjacent to Kotanitanga Bath (NS3B). Methylmercury levels were nearly an order of magnitude greater in Cub Bath (TS2) than in Tiger Bath (TS1), even though the total mercury levels were roughly the same (~1000 ng L^−1^).

**Table 2.** Mercury analyses of Ngawha Geothermal Field (NGF) hot springs: methylmercury (MeHg) and total mercury (HgT) and the fraction of total Hg as MeHg (as %) in filtered water samples, solid total mercury (THg(s)) in hot spring sediments, and gaseous elemental mercury (Hg(0)) above the springs. Temperature and pH measurements taken during each sampling campaign are provided as a range. Published reference and regional background values are included.

|  | T (°C)<br>Water | pH | MeHg<br>(ng L<sup>-1</sup>) | Hg<sub>T</sub><br>(ng L<sup>-1</sup>) | %MeHg<sub>T</sub><br>of Hg<sub>T</sub> | THg(s)<br>(µg g<sup>-1</sup>) | Hg(0)<br>(ng L<sup>-1</sup>) |
| --- | --- | --- | --- | --- | --- | --- | --- |
| **Ngawha Geothermal Field** |  |  |  |  |  |  |  |
| Tiger Bath (TS1) | 35.3–40.5 | 2.9–3.0 | 1.56 | 1031 | 0.15 | 3467 | 25 |
| Cub Bath (TS2) | 24.5–33.4 | 3.6–3.2 | 13.9 | 1042 | 1.33 | 1274 | 4.3 |
| Hole to right of Tiger Bath (TS3) | 35.9–40.3 | 2.9–3.8 | 4.33 | 16711 | 0.03 | 7000 | 7.5 |
| Hole 3 m from Tiger Bath (TS4) | 21.6 | 3.96 | 0.53 | 2160 | 0.02 | - | - |
| Cinnabar Bath (GN1) | 40.4 | 6.72 | 5.36 | 1461 | 0.37 | - | - |
| Jubilee Bath (GN2) | 38 | 6.45 | 1.28 | 244 | 0.52 | - | - |
| Twin Bath (GN3) | 34.1 | 6.44 | 1.61 | 722 | 0.22 | - | - |
| Kotanitanga (NS1B) | 32.4 | 6.54 | 4.28 | 487 | 0.88 | - | - |
| Adjacent to Favourite Bath (NS3B) | 26.5 | 7.14 | 3.57 | 253 | 1.41 | - | - |
| Waikato Bath (NS4B) | 45.5 | 6.7 | 0.90 | 326 | 0.28 | - | - |
| **Regional values** |  |  |  |  |  |  |  |
| NGF soils and waters(1) | - | - | - | 17000–710000 | - | 0.017–0.35 | 2000–78500 |
| Tiger Bath (1) | 32–57 | 5–8 | - | 710000 | - | - | - |
| NGF non-thermal waters (1) | - | - | - | 300–500 | - | - | - |
| Puhipuhi mine waters (2) | - | - | 0.61–1.02 | 69.6–240 | 0.3-1.6 | 35–1748 | 0.06–0.50 |
| Wairua River, Northland (3) | - | - | - | 100–1000 | - | <100 | - |
| **Yellowstone National Park** |  |  |  |  |  |  |  |
| Nymph Lake spring #1, YNP (4) | - | - | 0.026 | 170 | 0.02 | - | - |
| Nymph Lake spring #2, YNP (4) | - | - | 0.43 | 520 | 0.08 | - | - |
| Roadside West Spring, YNP (4) | 65 | 6.4 | 0.08 | 200 | 0.04 | - | - |
| YNP thermal waters and microbial mats (4) | 36.4–80 | 2.2–8.6 | <0.013–0.43 | 15-520 | - | 0.0049–120 | - |
| Acidic thermal springs YNP (5) | - | - | <0.025 | 38-94 | - | - | - |
| Acidophilic microbial mats, YNP (5) | - | - | 0.03-1.62 | 2-71 | - | 0.16-18 | - |
| YNP filtered thermal waters and microbial mats (6) | 21.9–74.4 | 1.8–9.3 | - | 7.3–144 | - | 0.38–20.52 | - |
| **Reference standards and values** |  |  |  |  |  |  |  |
| North American lakes (7,8) | - | - | 0.003–6 | 0.04–74 | 8.11 | - | - |
| North American river & streams (7,8) | - | - | 0.078–0.55 | 1–7 | 7.86 | - | - |
| NZ Drinking Water Standard | - | - | - | 7000 | - | - | - |
| US EPA, acute aquatic life water standard | - | - | - | 2400 | - | - | - |
| US EPA, chronic aquatic life water standard | - | - | - | 770 | - | - | - |

(1) Davey & Moort 1985, (2) Gionfriddo et al. 2014, (3) Hoggins & Brooks 1973, (4) King et al. 2006, (5) Boyd et al. 2009, (6) Wang et al. 2011, (7) Krabbenhoft et al. 2007, (8) USEPA(2001)

Sediments collected from the Tiger Springs area in October 2011 were analyzed for Hg_T_ (Table 2). Total Hg concentrations were highest in the hole adjacent to Tiger Bath (TS3) at ~7000 µg g^−1^. Tiger Bath (TS1) sediments had about half the solid Hg_T_ of TS3, at 3467 µg g^−1^, and Cub Bath sediments had about a third the solid Hg_T_ of Tiger Bath (1274 µg g^−1^). Gaseous Hg^0^ emissions were also recorded from the baths (Table 2). The Hg(0) concentration measured above Tiger Bath (25.0 ng L^−1^) was greater than that of Cub Bath (4.34 ng L^−1^), but also varied across the bath (4.38 – 25.0 ng L^−1^). Furthermore, these values are an order of magnitude lower than previously reported concentrations of fumarolic Hg(0) in the Tiger Springs area 13.5 – 276 µg L^−1^, and 710 µg L^−1^ from Tiger Bath specifically (32).

### Microbial diversity from metagenomic datasets

Phylogenetic analysis of ribosomal marker protein s3 from assembled Tiger and Cub Bath metagenomes identified members of Deltaproteobacteria, Gammaproteobacteria, Thermotogae, Micrarchaeota, Parvarchaeota, Thermoplasmata, and other Euryarchaeota (Figs. S4 and S5). Furthermore, phylogenetic analysis revealed a greater breadth in the phyla of genomes resolved from Cub Bath, compared to Tiger Bath, with Verrucomicrobia, Acidobacteria, Firmicutes, Planctomycetes, Alphaproteobacteria, and Betaproteobacteria identified in Cub Bath only (Fig. S4). Ribosomal proteins from Thermoprotei were identified only in assembled Tiger Bath metagenomic data (Fig. S5). Genome binning resulted in acidophilic and thermophilic bacteria and archaea dominating the Tiger and Cub Bath metagenomic datasets; with the average coverage of scaffolded contigs within genome bins ranging from 7.1 – 789x (Fig. 1; Table S2). The highest coverage bins in each bath were related to *Acidithiobacillus*, Thermotogales and *Thermoplasma* (Fig. 1).

**Figure 1.**
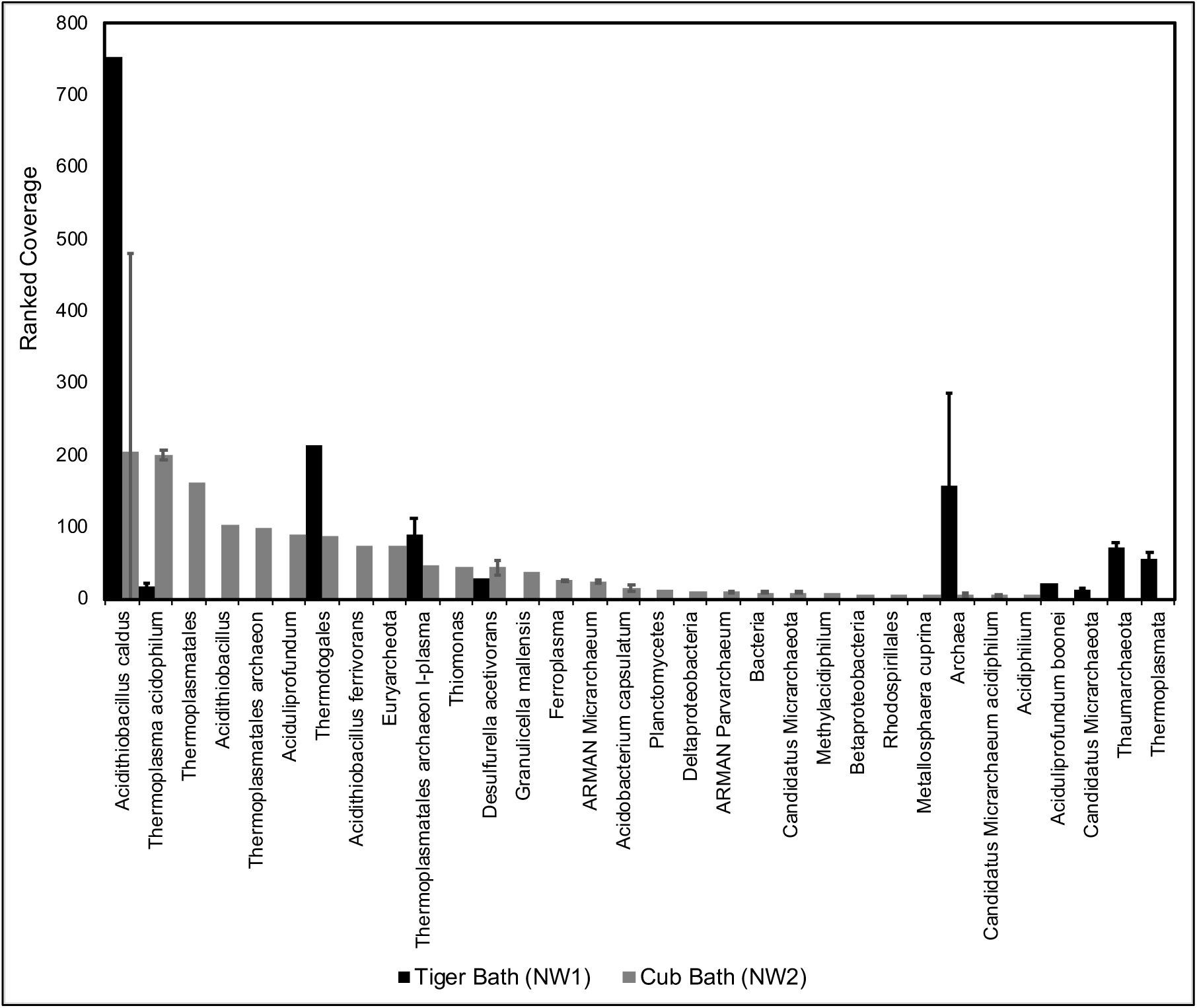
Rank abundance by scaffold coverage of ribosomal protein S3 within binned and unbinned genomes from A) Tiger (NW1) and B) Cub (NW2) bath metagenomes. Analyses and annotations performed in ggKbase (ggkbase.berkeley.edu/). Genomic bin phylogeny used for ribosomal S3 proteins from genomic bins, while scaffold phylogeny to the lowest common ancestor is given for unbinned ribosomal proteins. When multiple ribosomal protein S3 had the same taxonomic classification the average coverage is shown, with error bars representing standard deviation. Values ranked by NW2 coverage.

Genome bins were comprised of aerobes, as well as obligate and facultative anaerobes, capable of sulfur oxidation (*Acidithiobacillus* spp., *Thiomonas*), sulfur reduction (*Desulfurella acetivorans, Granulicella mallensis*), iron oxidation (*Ferroplasma*, *Acidithiobacillus ferrivorans)*, iron reduction (*Acidobacterium capsulatum*), methane oxidization (*Methylacidiphilum*), and acetate oxidation (*Desulfurella acetivorans*) (Fig. 2). To elucidate important biogeochemical links to Hg cycles mediated by these microbial phylotypes, metagenomes were searched for genes encoding for Hg, sulfur, sulfate, and methane cycling (Fig. 2). We note here that genes involved in methanogenesis (notably *mcrA*) and methane oxidation (*pmoA*) were nearly absent from metagenomes (Fig. 2).

**Figure 2.**
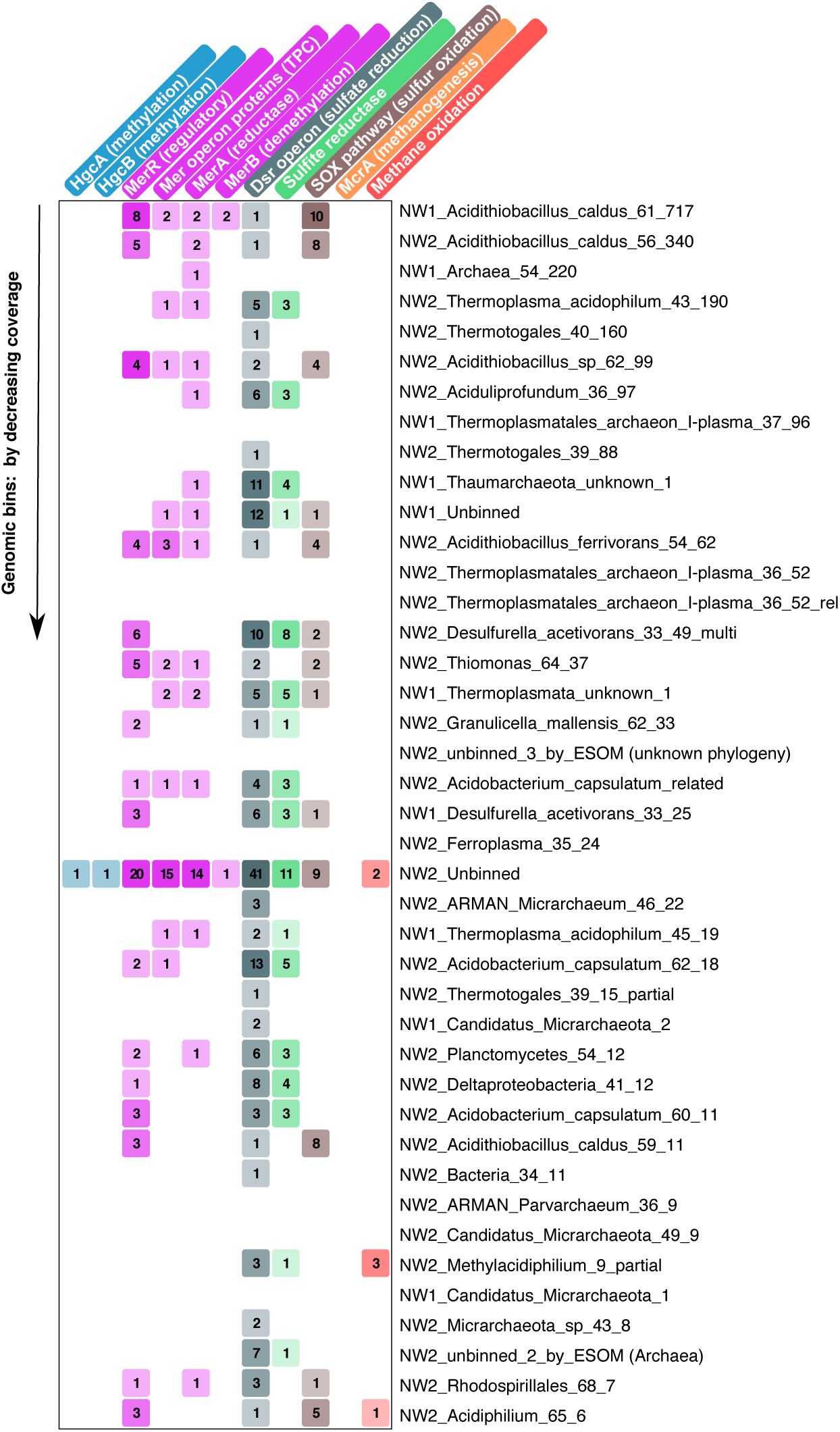
Heatmap showing functional proteins from various biogeochemical pathways associated with Hg within genomic bins and unbinned (NGAWHA_1_UNK, NGAWHA_2_UNK) scaffolds from Tiger (NW1) and Cub (NW2) bath metagenomes. Intensity of color refers to number of genes from each bin that encode for the enzyme, operon, or pathway involved in mercury, sulfur, or methane cycling; values provided in cell for reference. Genomic bins were ordered by the consensus coverage for all scaffolds within the bin (highest to lowest). Analyses and annotations were performed in ggKbase (ggkbase.berkeley.edu)

### Mer operon

Assembled metagenomes were screened for genes belonging to the *mer* operon that encode for mercuric reductase (*merA*), organomercurial lyase (*merB*), a periplasmic protein (*merP*), and inner membrane proteins involved in Hg(II) transport (*merT*, *merC*, *merE*, *merF*, *merG*), as well as one or more regulatory proteins (*merR*, *merD*) (13). At Ngawha, scaffolded *mer* genes were often encoded in the higher coverage genome bins of each bath, *Acidithiobacillus* spp., *Thiomonas*, and Thermotogae (Table S5; Fig. 2). The coverage of *mer* scaffolds ranged from 3.9 – 836x, with scaffold lengths of 1,021 – 101,352 bp, indicating that a significant number of reads mapped to each sequence. A high fraction of reads (3.24E-04 – 1.94E-04, Table S1) from each metagenome were predicted to encode mercuric reductase (MerA) using the HMM-model. BlastP analysis of HMM-search outputs indicated that the HMM-model was insufficient for filtering sequences that are predicted to encode for MerA paralogs dihydrolipoamide dehydrogenase and pyridine nucleotide-disulfide oxidoreductase. Therefore, the HMM-model likely over-estimates MerA abundance encoded by the raw reads. MerA homologues identified in the assembled Tiger and Cub Bath metagenomes using ggKbase annotations were related to both Archaea (Euryarchaeota), and Bacteria (Proteobacteria and Bacteroidetes) (Fig. 3). Most archaeal MerA were related to *Thermoplasma*; however, four MerA homologues branched deeply from known archaeal homologues and were related (<43%, BlastP sequence alignment) to MerA from *Candidatus* Methanoperedens nitroreducens (NW1 scaffolds 675 and 247, and NW2 scaffolds 1319 and 13696; Fig. 3). Most MerA homologues from Cub Bath were related to *Acidithiobacillus* spp. (Fig. 3). MerA homologues identified in the metagenomes were from operons also encoding for MerP, MerT, and MerR (Table S5). Several scaffolds related to Euryarchaeota (*Thermoplasma*) encoded just for MerA and MerP (NW1_scaffold_26, NW1_scaffold_675, NW2_scaffold_876; Table S5), consistent with *Thermoplasma mer* genes sequenced from acid mine drainage (AMD) (52).

**Figure 3.**
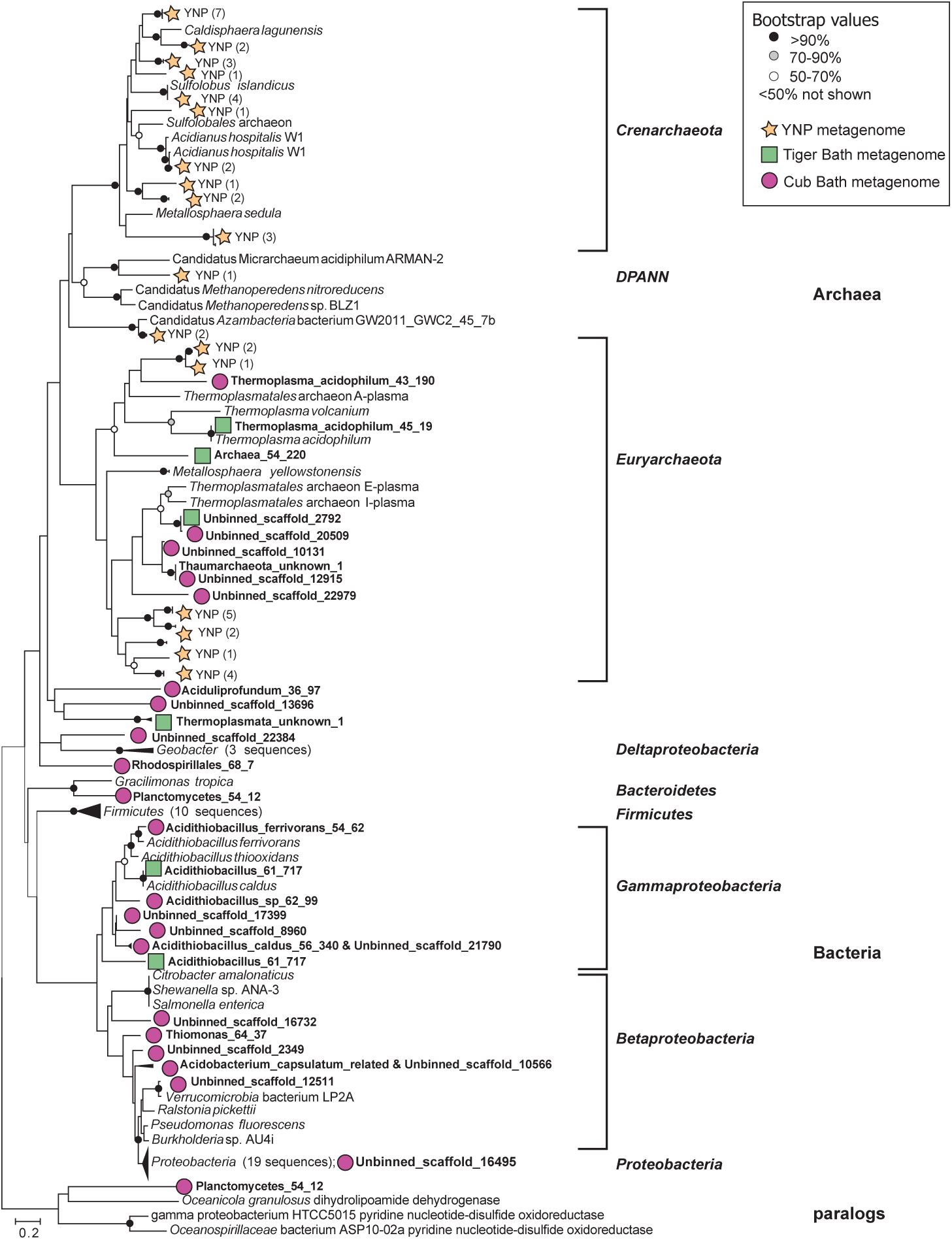
Maximum likelihood tree showing MerA phylogeny of 31 sequences pulled from assembled Tiger (NW1) and Cub (NW2) Bath metagenomes using ggKbase. Genomic bins or scaffold ID (when MerA was unbinned) are given in bold. Included in analysis are 113 MerA homologues, including 46 sequences from Yellowstone National Park (YNP) metagenomes (Inskeep et al. 2013). Trees were constructed using Le Gascuel amino-acid substitution model with gamma distribution in MEGA6 (Tamura et al. 2013). Bootstrapped with 100 replications. The initial Neighbor-Joining tree was constructed with pairwise distances estimated using a JTT model. Positions with less than 90% site coverage (e.g. alignment gaps, missing data or ambiguous bases) were excluded. A total of 419 positions were used in the final dataset.

Two similar but distinct *merB* genes were identified in assembled Tiger and Cub Bath metagenomes, related to *Acidithiobacillus caldus* and *Thioalkavibrio* spp., respectively (Fig. S6). The *merB* gene from Tiger Bath (NW1_scaffold_113) was linked to a genome bin identified as *Acidithiobacillus caldus*, with an average scaffold coverage of 750x (Table S5). The same scaffold (NW1_scaffold_113) contained two *merR* genes that were convergent and divergent, respectively to *mer* genes that encode for MerT, MerP, and MerA. The *merB* gene from Cub Bath (NW2_scaffold_17399) was most closely related to *merB* from *Thioalkavibrio* spp.; however the scaffold itself was unbinned, with an overall taxonomic identification of *Acidithiobacillus ferrivorans* (Table S5). The scaffold also contained *merR* and *merA* genes related to *Acidithiobacillus ferrivorans* (WP_035195121). The translated organomercurial lyases (MerB) from Tiger and Cub Baths aligned to conserved cysteines(53), indicative of their true functionality (Fig S7).

### Biological Hg methylation

Both Tiger and Cub Bath metagenomic read sets were searched for sequences sharing homology to Hg methylation genes (*hgcA* and *hgcB*) (17). A small fraction of reads (2.16E-07) were identified as fragments of *hgcA* sequences in the Cub Bath metagenome (Table S1). The nucleotide reads were each 100 bp in length, and when translated, aligned to several regions of HgcA from known methylators, including the highly conserved “G(I/V)NVWC” region of the HgcA protein ((17), Fig. S8). Several of the reads aligned to one another, and a composite amino acid sequence (59 AA in length) is shown in Fig. S8. Phylogenetic analyses of the translated reads revealed two distinct *hgcA*-like genes in the Cub Bath metagenome (Fig. 4). One of the reads closely matched (BlastP search) to pterin-binding regions of HgcA-like proteins from marine bacteria *Streptomyces* sp. CNQ-509 and *Nitrospina* spp. (E-value: 8e-05; 93% sequence coverage; 55%), and to fused HgcAB proteins primarily found in thermophilic archaeal and bacterial genomes, *Thermococcus* sp. EP1, *Kosmotoga pacific*, *Pyrococcus furiosis*, and *Methanococcoides methylutens* (E-value: 3E-06 – 8E-08; 100% sequence coverage; 65–75% sequence ID). Whether these microbes with genomes encoding for an HgcAB fused protein are capable of Hg methylation is unknown (16). When tested for Hg methylation capability, both *Pyrococcus furiosus* and *Methanococcoides methylutens* were unable to produce MeHg above controls (16, 54).

**Figure 4.**
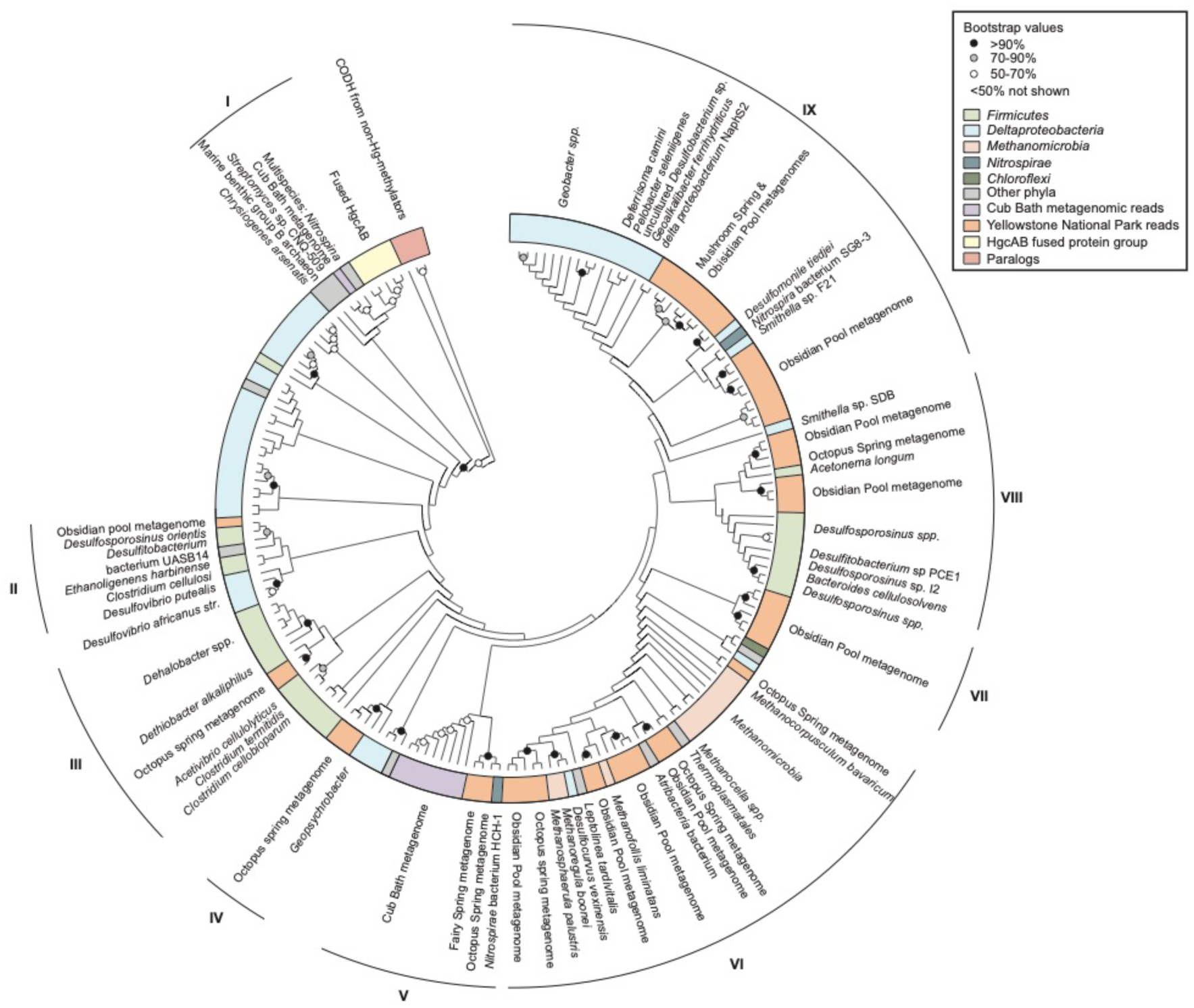
Maximum likelihood tree showing HgcA phylogeny of reads obtained from the Cub Bath (NW2) metagenome using HMM-search. Included in analysis are 183 sequences, including 56 HgcA homologues from YNP metagenomes, pulled from JGI (Table S9). Amino acid sequences are compared to HgcA homologues from known and predicted methylators. Also included are HgcAB fused proteins from hyperthermophilic bacteria and archaea, and carbon monoxide dehydrogenase/acetyl-CoA synthase subunit gamma (HgcA paralogs) from non-methylators. Trees were inferred from the Le Gascuel amino-acid substitution model with gamma distribution in MEGA6 (Tamura et al. 2013). Bootstrapped with 100 replications. The initial Neighbor-Joining tree was constructed with pairwise distances estimated using a JTT model. Positions with less than 93% site coverage (e.g. alignment gaps, missing data or ambiguous bases) were excluded. A total of 55 positions were used in the final dataset. Groups (I-IX) designate distinct subtrees that contain HgcA from hot spring metagenomes. Representative reference sequences are labeled within each subtree, while phylogeny of all branches are indicated by color.

The second set of *hgcA* reads from Cub Bath (reads 2–7; Fig. 4), including the composite sequence, aligned using BlastP to the pterin-binding region of HgcA proteins from known and predicted Hg methylators *Desulfosporosinus youngiae*, *Clostridium cellobioparum*, *Desulfosporosinus* sp. Tol-M, *Desulfosporosinus* sp. BRH-c37(E-value: 4E-21 to 7E-22; 100% query cover; 71–75% ID). Phylogenetic analyses (Fig. 4) of the composite sequence revealed homology to HgcA from *Nitrospirae* bacterium HCH-1 (Genbank ID: LNQR00000000.1) and to HgcA from hot spring metagenomes. The *hgcA* sequences pulled from assembled YNP and British Columbia hot spring metagenomes from JGI-IMG databases span a wide range of diverse phyla and are predicted to encode HgcA that relate closely to those from Deltaproteobacteria, Firmicutes, Nitrospirae, Planctomycetes, and Euryarchaeota (Methanomicrobia and Thermoplasmata) (Fig. 4). BLASTP searches of translated amino acid sequences revealed common closest matches (<83% ID) to *Clostridium straminisolvens*, *Deltaproteobacterium* sp. NaphS2, *Nitrospirae* bacteria SG8-3 and HCH-1, *Physcisphaerae* bacterium SG8-4, *Smithella* spp., *Methanocella arvoryzae*, and *Thermoplasmatales* archaeon DG-70-1. Of these closest relatives, only *Thermoplasmatales* archaeon DG-70-1 is from a thermophilic phylum (Thermoplasmata); however, this strain was isolated from an anaerobic, moderately halophilic, and mesophilic aquatic environment (55), rather than a geothermal spring.

A high fraction of reads from each metagenome aligned to *hgcB* sequences from known and predicted methylators (2.08E-06 – 3.02E-06; Table S1). However, BLAST analysis of the translated *hgcB*-like genes resulted in closest matches to ferredoxin-encoding gene sequences in both Tiger and Cub Bath. To differentiate between other ferredoxin proteins and HgcB homologues, the translated *hgcB*-like genes were searched for the conserved “C(M/I)ECGAC” site in HgcB (17). A complete *hgcB* gene was identified in the assembled Cub Bath metagenome (NW2_scaffold_10600_2, Supp. Mat.). A translated BlastP search of the sequence revealed closest matches (E-value: 2E-26–3E-29; 91–98% coverage; 53– 57% identity) to HgcB from predicted Hg methylators *Dehalobacter* spp., *Clostridium straminisolvens*, *Nitrospirae* bacterium HCH-1, and *Syntrophobotulus glycolicus*. Other features on the assembled contig that contains the *hgcB* gene (NW2_idba_contig_10600), include a partial *hgcA* gene upstream from *hgcB*, and downstream genes encode for a DsrE/DsrF-like family protein (involved in intracellular sulfur reduction) related to Planctomycetes; a copper-binding protein related to Aquificae; and a putative transcriptional regulator related to ArsR repressor, and an arsenical pump membrane protein (ArsB), both related to Firmicutes (Fig. S9).

### Biological sulfur species cycling in Tiger and Cub Baths

The availability and speciation of sulfur compounds within geothermal environments constrains the strategies employed by microorganisms to deal with Hg in these ecosystems. For example, Hg methylation is impeded in ecosystems with elevated aqueous sulfide concentrations (56). Conversely, the availability of soluble Hg to microorganisms is limited by the solubility and/or microbially-mediated dissolution of cinnabar/metacinnabar (57). To elucidate these interactions, the Cub and Tiger Bath metagenomes were searched for genes related to dissimilatory sulfite/sulfate reduction using *dsrAB* genes that encode subunits A and B of the dissimilatory bi(sulfite) reductase enzyme (DsrAB); (58), and the *sox* pathway genes (*soxR*, *soxXA, soxYZ*, *soxB, sox(CD)*_2_, *soxG*) that encode enzymes used in hydrogen sulfide and elemental sulfur oxidation (59).

Complete or near complete *dsrAB* sequences were recovered from assembled Tiger and Cub Bath metagenomes using ggKbase annotations (Table S6). The genes were often present on scaffolds containing other genes encoding proteins involved in sulfite/sulfate reduction to sulfide. Both *dsrC* and *dsrD* genes were present on scaffolds with *dsrAB* in NW1_Thaumarchaeota_unknown_1, NW2_Deltaproteobacteria_41_12, and NW2_Acidobacterium_capsulatum_related. Phylogenetic analyses of the translated genes indicated six distinct groups of DsrAB sequences in Tiger and Cub Baths (Fig. 5, Table S6), Groups I-VI. Three distinct groups of reductive bacterial type DsrAB groups (I-III) were identified, most closely related to known thermophilic sulfur-reducing bacteria such as *Desulfurella acetivorans* as well as uncultured *Gemmatimonas* sp. Sg8-17. Groups IV and V contained sequences related to reductive archaeal DsrAB in genomic bins related to *Thermoplasmata*, *Thermoplasma acidophilum*, and *Aciduliprofundum*. Group VI DsrAB contain sequences distantly related to *Candidatus* Rokubacteria CSP1-6, *Caldivirga maquilingensis* and *Thermodesulfobacterium* spp. (WP_051754629). Group V DsrAB were likely uncultured reductive archaeal type DsrAB related to *Vulcanisaeta*; unbinned sequences were from scaffolds with a majority of sequences related to Archaea. Group VI DsrAB most closely related (>50%) to DsrAB from the sulfate-reducer *Candidatus* Rokubacteria CSP1-6 (60); the sequence was distinct to DsrA from Group IV and is from genomic bin NW2_Planctomycetes_54_12 (Table S6). The DsrAB phylogenies as represented in Fig. 5 agree with the phylogenetic annotations of the scaffolds from which the genes were obtained (see Table S6). Importantly, while there is evidence for bacterial and archaeal sulfate and sulfur reduction within Tiger and Cub Baths, none of the DsrAB groups detected (Groups I-VI, Fig. 5) contain known Hg methylators from the primary sulfate-reducing phyla, Deltaproteobacteria and Firmicutes (18). Nor were any *hgcAB* genes present in sulfate-reducing genomic bins from Tiger or Cub Bath.

**Figure 5.**
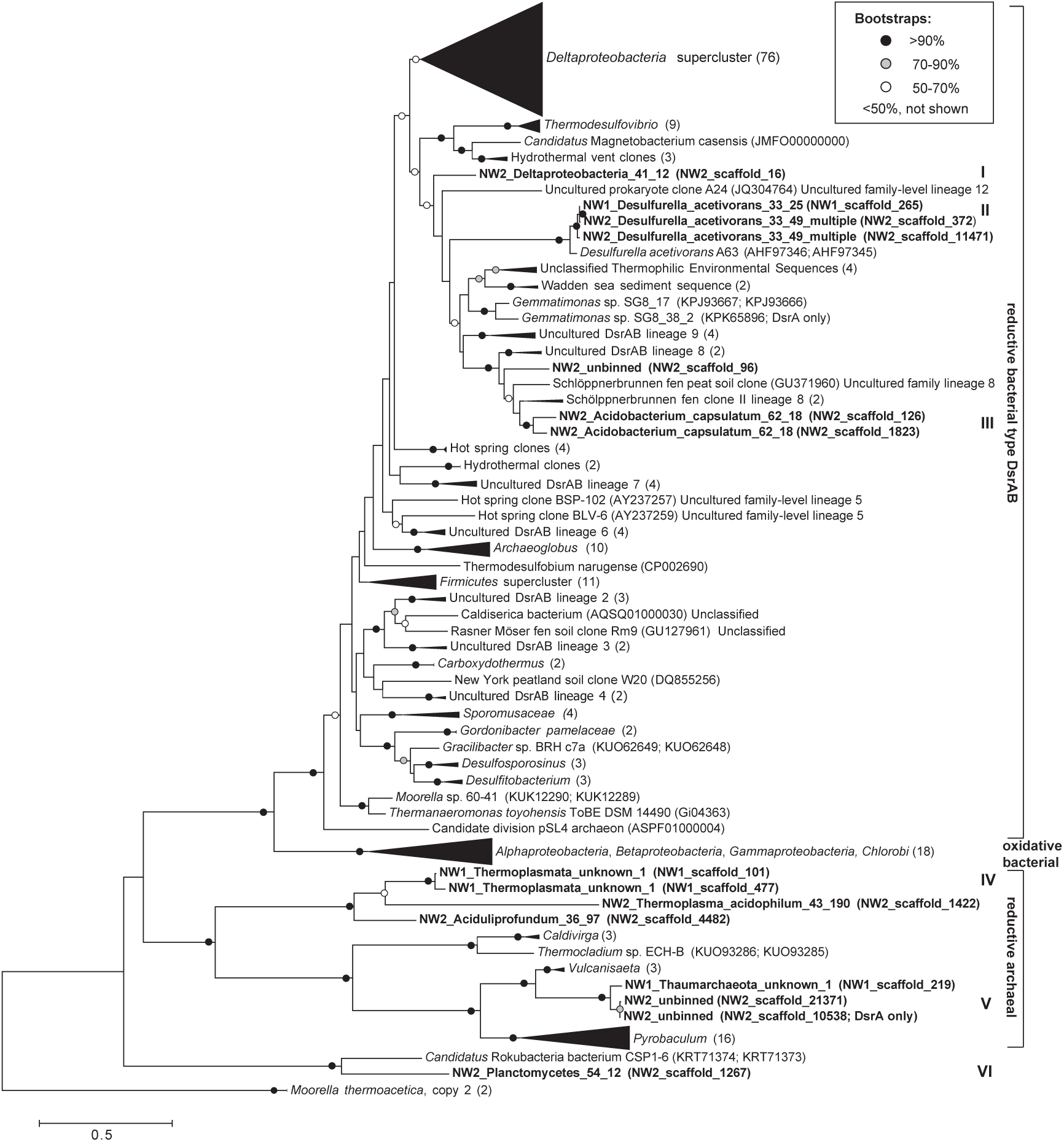
Phylogenetic analysis of DsrAB by Maximum Likelihood method. The 15 DsrAB homologues in this study were pulled from idba-UD assembled Tiger and Cub metagenomes using ggKbase. They were compared to 218 reference DsrAB sequences pulled from the Dome database, including 15 representative DsrAB sequences from thermophilic environments (Muller et al. 2015). There were six distinct groups of DsrAB homologues found at Ngawha. The tree was constructed using MEGA 6 (Tamura et al. 2013) with Le Gascuel 2008 model with gamma distribution, pairwise distances estimated using a JTT model. All positions with less than 95% site coverage were eliminated, including alignment gaps, missing data, and ambiguous bases. There were a total of 492 positions in the final dataset, with 100 bootstrap replicates.

Genes encoding for sulfur-oxidation *(sox*) in acidophilic Beta- and Gammaproteobacteria were detected in both Tiger and Cub Bath metagenomes and were related to *Acidithiobacillus* spp. and *Thiomonas* spp. (Fig. 2; Table S7). Furthermore, sequences from archaeal and bacterial *sox* pathways, including from acidophiles *Acidiphilium* and *Acidocella*, were present in the unbinned metagenomic data (Table S7).

## DISCUSSION

Geothermal systems provide an environment in which relationships between the chemical and physical processes controlling Hg speciation and bioavailability, and microbial Hg transformations (4, 7), remain poorly understood. Temperature and pH constitute major drivers of microbial diversity in geothermal springs, with pH contributing to a greater extent (61, 62). Indeed, previous studies show that acidic geothermal spring communities appear quite distinct from those of neutral and alkali springs, irrespective of temperature (61, 62). In our study, despite their mutual close proximity and broadly similar physicochemical properties and dissolved total Hg concentrations, the Tiger and Cub Baths of the NGF hosted very distinct microbiomes (Fig. S10). The greater diversity in genomic bins representative of the Cub Bath microbiome, compared to Tiger Bath (Fig. 1), may have promoted Fe and S redox cycling to a greater extent, which in turn could facilitate the dissolution of metal sulfides such as pyrite or cinnabar. Subsequently, this dissolution could have increased the bioavailability of Hg(II) for *hgcAB*+ equipped microorganisms, by oxidizing reduced S and increasing dissolved Hg(II), respectively (Fig. 6).

**Figure 6.**
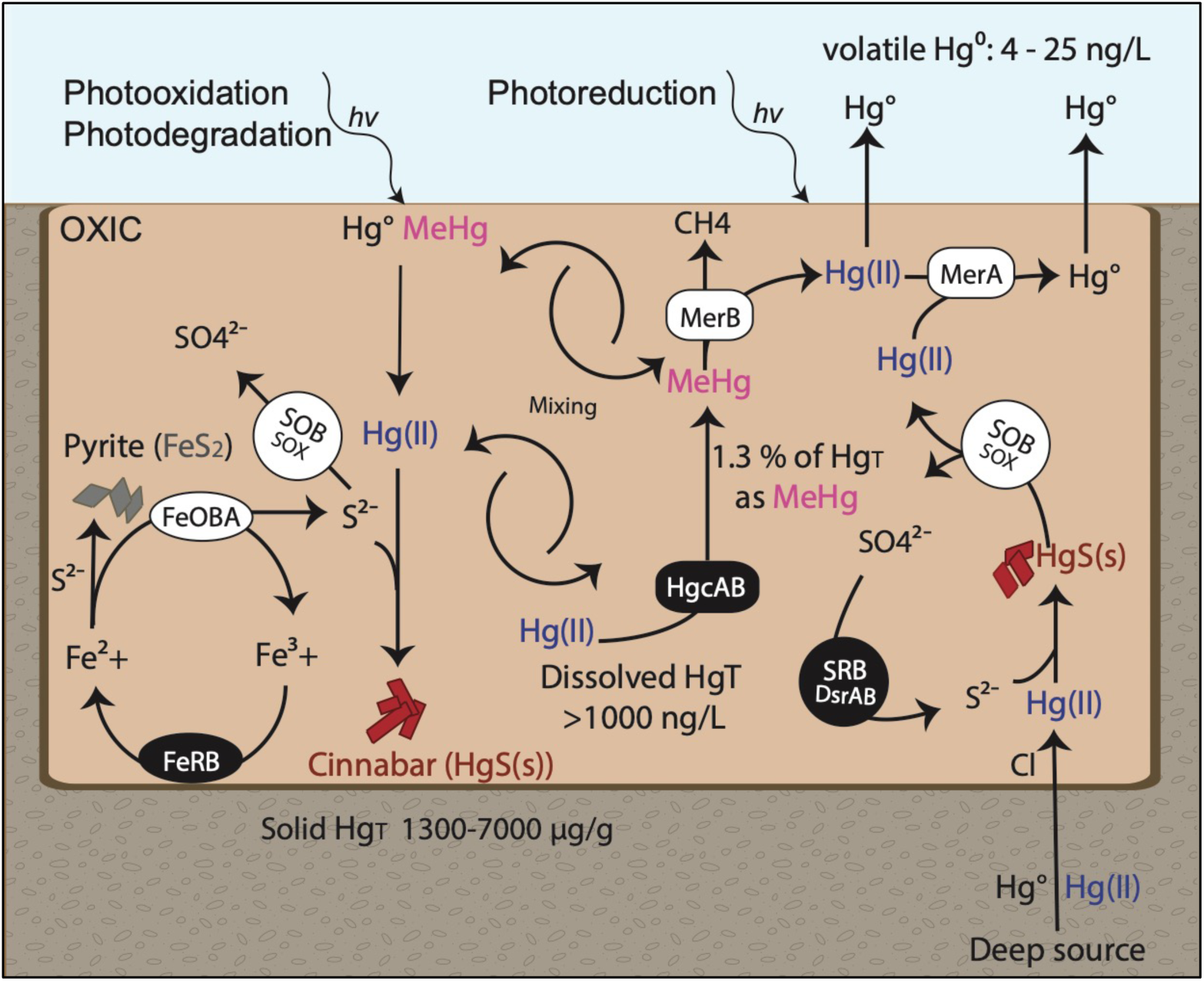
Conceptual model of biogeochemical cycling of mercury (Hg), sulfur (S), and iron (Fe) in Hg-enriched, sulfidic, low pH mesothermal springs. Gaseous elemental mercury (Hg0) (as well as Hg(II)) from deep geological sources enters the surface waters of the springs where it becomes oxidized to Hg(II) (enhanced by chloride (Cl))(32) and then complexes with sulfides (S^2−^) to produce cinnabar (red rhombohedral symbols; HgS(s)). Round icons represent microbial-mediated reactions, white are primarily aerobic-associated mechanisms, and black are primarily anaerobic. Sulfur-oxidizing bacteria (SOB) equipped with the SOX-pathway along with Fe-oxidizing bacteria and archaea (FeOBA) are able to enhance dissolution of metal sulfides, such as pyrite (silver rhombohedral symbols; FeS2) and HgS(s). Sulfate-reducing bacteria (SRB) and Fe-reducing bacteria (FeRB) further mediate the redox chemistry of S, Fe, and Hg. As Hg(II) becomes bioavailable to microbes it can be reduced to Hg(0) by microbes equipped with mercuric reductase (MerA) or methylated to MeHg by HgcAB-equipped microbes. MeHg can be demethylated by MerB-equipped microbes to Hg(II) and CH4, and then reduced to Hg(0) by MerA. At the surface of the springs, photoreduction can also contribute to Hg(II) reduction to Hg(0), as well as the degradation of MeHg. Photolytic oxidation may also counter Hg(0) volatilization from surface waters, keeping Hg(II) in spring water to be transformed by microbes or partitioned to sulfide minerals. Advective mixing of spring waters ensures that Hg species travel across the redox boundaries that likely partition the Mer-equipped microbes to oxic surface waters from the HgcAB-equipped microbes that likely occupy the anaerobic sediment/water boundary.

NGF genomes featured similar metabolic capabilities to those recovered from AMD. The oxidation of reduced sulfur species by both aerobic and anaerobic chemolithotrophic microorganisms produces electrons utilized in respiration and CO_2_ assimilation (63), critical processes for living in AMD and/or geothermal systems (63, 64). Dominant members of the Tiger Springs’ communities, *Acidithiobacillus spp.* and *Thiomonas spp.*, can utilize various metal and sulfur-oxidizing enzymes, pathways, electron transport mechanisms, and substrates (63) to sustain activity. *Acidithiobacillus ferrivorans* is an obligate chemolithoautotroph and facultative anaerobe that oxidizes Fe(II); some strains are also able to utilize sulfur, thiosulfate, tetrathionate, and pyrite (65). *Acidithiobacillus ferrooxidans* can utilize metal sulfides to support growth (63), and *Acidithiobacillus caldus* is also capable of oxidizing reduced inorganic sulfur species, producing sulfate via the *sox* pathway (66). A number of other NGF genome bins, including several associated with *Thermoplasma* and *Thiomonas* spp., were equipped to respire using either sulfur or organic carbon (30), and *Thiomonas* can also oxidize arsenite (As(III)) to arsenate (As(V)) (67). The *Thiomonas-*like genome bin from Cub Bath showed evidence for the presence of both *sox* and arsenite oxidase pathways (Figure S2).

The microbiomes of Tiger Springs were also similar to those found in AMD with respect to stress resistance/response mechanisms for acid and heavy metals (e.g., (30, 68)), featuring mechanisms for pH homeostasis, for example. The highest genome coverage in each bath was associated with *Acidithiobacillus caldus*, a microorganism equipped with multiple heavy metal resistance pathways (*ars*, *mer*, *czc*, and Te^R^). The *mer* genes in Tiger Spring metagenomes were predominately found in aerobic bacterial and archaeal genome bins (Fig. 2; Table S5), particularly the mesophilic acidophiles *Thermoplasma* and *Acidithiobacillus* (Fig. 3). This finding was in sharp contrast to the diversity of *merA* genes in YNP metagenomic datasets, which were principally from archaeal taxa (e.g. Sulfolobales, Acidobales, DPANN). Based on the observations of Geesey et al. (4) that archaea dominated acid springs with high Hg content, and given that MerA homologues are often encoded in the genomes of acidophilic archaea (4, 12, 13), we expected that archaeal MerA would dominate in Tiger and Cub Baths.

While Tiger and Cub Baths are considered acidic (pH <4), they are substantially lower in temperature than springs studied in YNP and the Western US; these different observations may be due to a number of factors. First, Geesey et al. (4) noted that the number of bacterial MerA homologues detected in acid springs increases with decreasing temperature (from >73 °C to <55°C). Thus, the lower temperatures at NGF may explain the presence of a larger number of bacterial MerA homologues. Second, bacterial taxa are also known to be rare in higher temperature (>65 °C) acid (<pH 4) geothermal ecosystems (69, 70). Thus, the minimal distribution of bacteria with *mer* genes was most likely a function of low thermophilic acidophile diversity, rather than the absence of a taxonomic capacity to transform Hg.

A notable feature of Cub Bath was the higher percentage of Hg_T_ present as MeHg_T_, compared to Tiger Bath. We speculate that this finding reflects a greater degree of microbially-mediated turnover of aqueous Hg(II) to MeHg in the former spring. However, the difference in MeHg levels between the baths could also be indicative of higher demethylation rates in Tiger Bath compared to Cub Bath. While *merB* genes were identified in both baths, the sequencing coverage of *merB* scaffolds was higher in Tiger Bath (836x) compared to Cub Bath (18x) (Table S5). While abiotic methylation in geothermal waters is not yet well understood (71), such processes are not likely to account for the higher MeHg levels in Cub Bath relative to Tiger Bath. Furthermore, NGF MeHg values were 1-2 orders of magnitude greater than values recorded at some YNP hot springs (5, 6). An analysis of NGF metagenomes found that *hgcA* and *hgcB* genes were only detected in Cub Bath and not Tiger Bath. Cub Bath Hg methylation genes are most likely bacterial, belonging to a sulfate-reducing clade closely related to Firmicutes and Deltaproteobacteria, possibly from the Nitrospirae (Fig. S9). Microbial Hg-methylation via active sulfate-reducing microorganisms is consistent with the greater observed amount of MeHg (~1.3% of dissolved total Hg) alongside higher concentrations of reduced sulfur (3.5x) in Cub Bath. Of particular note was an *hgcAB*+ sulfate-reducing bacterium in Cub Bath metagenome (NW2_unbinned) that may represent a novel acidophilic Hg methylator equipped with heavy metal resistance. This genome encoded *hgcA* and *hgcB* on a scaffold with a putative copper chaperone (HMA/CopZ), arsenical resistance operon repressor (ArsR), and arsenical pump membrane protein (ArsB) (Fig. S9). There are few reports of arsenate reduction by sulfate-reducing bacteria (72), although putative *ars* genes have been found in several species of *Desulfovibrio*, *Desulfosporosinus*, *Desulfomicrobium*, and *Desulfotalea*. Genomes of known, predicted, and non-Hg-methylating bacteria and archaea were searched for homologous proteins to those encoded on NW2_scaffold_10600 (Table S8). Notably, ArsR family transcriptional regulators are encoded directly upstream to HgcA in *Desulfovibrio desulfuricans* ND132 (DND132_1054) and *Desulfomicrobium baculatum* (Dbac_0377). While homologous ArsR proteins are common in genomes containing HgcA (Table S8), no genomes encode all of these proteins, although several genomes of known and predicted methylators encode for copper chaperones and arsenic resistance proteins, providing a measure of confidence in genome binning results. Notably, three homologous proteins are encoded in the genomes of *Desulfosporosinus* spp., often found in sulfate-rich, heavy metal-contaminated, low pH environments (73), and *Desulfosporosinus acidiphilus*, with an optimum growth pH of 3.6 – 5.5, was the first acidophile observed to methylate Hg (18). Therefore, we infer that the Cub Bath bacterium is likely a similar taxon capable of both Hg-methylation and As(V) reduction.

One of the strongest influences on Hg speciation and bioavailability (for methylation or volatilization) in acidic sulfidic hot springs is the formation and dissolution of cinnabar (HgS_(s)_), which in turn is impacted by the activity of sulfur- and iron-oxidizing microorganisms (5, 32, 56, 74, 75). A previous study (32) identified HgS_(s)_ as the most abundant and widespread Hg-bearing mineral in the Ngawha region, an observation we confirmed by XRD analysis from a topsoil sample in the Tiger Springs area (Fig. S2). The observed physicochemical conditions in Tiger and Cub Baths were on the cusp of HgS_(s)_ formation/dissolution, at pH 2-3 and pH > 4 (76). Thus the bioavailability of Hg(II) in the acid warm springs of the NGF reflected Hg solubility in the context of sulfur speciation and acidic conditions (56, 57). In aerobic sediments, sulfur-and iron-oxidizing bacteria and archaea may enhance the dissolution of cinnabar and increase Hg(II) bioavailability. Like sulfide, chloride can also affect Hg bioavailability to methylating microorganisms; methylation rates have been shown to correlate inversely with increasing chloride (Cl^−^) concentration (77, 78). Chloride concentrations in Cub Bath were higher than those of typical freshwaters (0.82 mM), and in combination with low pH, could have decreased net Hg methylation rates without impacting the viability of methylating microorganisms (77). Other factors that may influence Hg bioavailability include dissolved organic material (DOM) and thiol-DOM interactions (79–81), and turbidity which likely limits photolytic Hg transformations.

Together, these findings can account for the nearly 10x amount of filtered MeHg concentrations in Cub Bath, from which *hgcAB*+ equipped genome bins were recovered, compared to Tiger Bath, despite their near identical Hg_T_ concentrations in filtered water samples and the nearly 3x higher solid Hg content of Tiger Bath. In contrast, no *hgcA* reads were detected in the Tiger Bath metagenome at the sequencing depth of our study. Thus, bioavailable Hg may be getting enzymatically reduced or re-complexed by Hg-binding ligands (e.g., Cl^−^, DOM, S^2−^). Indeed, the low flux of Hg(0) from the surface of the bath suggests that most of the microbially-reduced Hg(II) remains dissolved, and potentially is continuously cycled between Hg(0) and Hg(II) redox states. Advective mixing of anoxic and oxic waters would also promote Hg cycling and exchange between aerobes and anaerobes, and likewise *mer*-equipped and *hgcAB*-equipped microbes, in the acid warm springs (Fig. 6).

While acidophilic microbial mats in YNP have been shown to accumulate MeHg and may actively methylate Hg (5, 6), the relative difference in pH between the two NGF springs is minimal, and therefore unlikely to account for the difference in recovery of *hgcAB* genes. An alternative explanation may be found in the Hg_(T)_ concentration in the sediments of both springs. The total solid Hg concentration in Tiger Bath was nearly three times higher than that of Cub Bath (3467 µg g^−1^ and 1274 µg g^−1^ respectively). Similarly, a small nascent hot spring located adjacent to Tiger Bath (TS3) had extremely high concentrations of both total dissolved and solid Hg (16,700 ng L^−1^ and 7000 µg g^−1^ respectively). Taken together, these observations suggest that the area immediately adjacent to Tiger Bath is a highly localized Hg “hotspot”, with elevated Hg levels selecting for microbial genomes encoding for Hg resistance. Hg methylation capability apparently does not confer Hg(II) resistance (82), and with the exception of two *Geobacter* spp., *mer* operon genes appear to be absent in genomes of known *hgcAB*^+^ microorganisms (14, 27). These observations are further corroborated by spike-in experiments in river water sediments that showed microbial Hg methylation was inhibited by Hg concentrations as low as 15.3 µg g^−1^ (24). Another plausible control on Hg methylation in the two baths may have been temperature, which was recorded as nearly 10°C different between baths during each sampling season (Table 2). Similar differences in temperature were recorded in the sediments (Tiger Bath at 51.1– 68.6°C and Cub Bath at 43.5–60.3°C). Principal component analysis of covariance between geochemical parameters measured in all springs across the NGF (Tables 1 and 2) indicates an inverse relationship between MeHg and spring temperature (Figure S11). Previous work shows that known *hgcAB+* methylators and predicted methylators are almost exclusively mesophiles with growth optima ~30°C (18). Known exceptions include the thermophile and known Hg-methylator *Desulfacinum hydrothermale* (83), psychrotolerant and psychrophilic predicted methylators *Geopsychrobacter electrodiphilus* and *Methanolobus psychrophilus* R15, and a group of hyperthermophiles possessing a fused *hgcAB*-like gene of unknown methylation functionality (16).

### Conceptual model of Hg speciation in warm acidic hot springs

In the NGF acid warm springs, Hg biogeochemical cycling reflects the detected or inferred microbially-mediated Hg transformations, as well as constraints imposed by metal sulfide solubility and vigorous microbially-mediated reduce S- and metal-oxidation reactions. The main abiotic and biotic controls on Hg cycling in acidic mesothermal springs are depicted in the conceptual model shown in Fig. 6. The bioavailability of Hg to Mer- and HgcAB-equipped microorganisms is controlled by cinnabar precipitation and dissolution, which in turn is influenced by pH and the presence of sulfur- and iron-oxidizing and sulfate-and iron-reducing bacteria and archaea. Bioavailable Hg(II) is methylated to MeHg by microbes, putatively sulfate-reducing bacteria, equipped with HgcAB. This process is limited by demethylation of MeHg (via MerB) and reduction (via MerA) of Hg(II) to Hg(0), by facultative anaerobic and aerobic iron-cycling and sulfur-oxidizing microorganisms. The volatile Hg(0) may evolve from spring waters, or be photo-oxidized and recycled to Hg(II). A significant sink for Hg(II) within the springs involves formation of solid HgS (metacinnabar, cinnabar), and potentially the adsorption of MeHg to sediments or particulate organic matter (which was not assessed here). However, in the acidic conditions, most MeHg and Hg(II) should remain dissolved, and could be continuously cycled between methylating and demethylating microorganisms. Importantly, temperature (>50 °C) and elevated Hg_(T)_ concentrations will restrict microbial methylation of bioavailable Hg(II). However, as geothermal inputs mix with cooler water, the microbial Hg methylation potential increases. Therefore, surface waters and groundwaters that receive geothermal inputs, such as catchment waterways down-gradient from NGF springs and discharges from hydrothermal power plants, may be important environmental pathways for MeHg mobilization and bioaccumulation.

